# Decoding peripheral stimulation from cortical and spinal recordings reveals complementary sensorimotor information

**DOI:** 10.64898/2026.05.05.722715

**Authors:** Laura Toni, Giulia Caffi, Ausilia Mazzarino, Birgit Nierula, Daniele Emedoli, Filippo Agnesi, Sandro Iannaccone, Falk Eippert, Fiorenzo Artoni, Silvestro Micera

**Author notes:** These authors contributed equally (Laura Toni, Giulia Caffi). These authors contributed equally (Fiorenzo Artoni, Silvestro Micera). Corresponding author: Laura Toni.

## Abstract

The somatosensory system encodes peripheral inputs through a sequence of ascending neural relays spanning spinal, subcortical and cortical levels. While multivariate decoding of electroencephalography (EEG) signals has demonstrated that cortical activity contains fine-grained information about somatosensory stimuli, the extent to which earlier processing stages contribute additional, non-redundant information remains unclear. To address this gap, we investigated a dataset comprising peripheral sensory stimulation and mixed stimulation (i.e., stimulation engaging both sensory and motor fibers). We assessed whether stimulation characteristics can be decoded from spinal recordings using high-density electrospinography (ESG), and whether combining ESG with EEG enhances decoding performance. Decoding accuracy varied systematically with both stimulation type and signal modality. ESG was the most informative signal for mixed and mixed vs sensory discrimination, reaching an average accuracy of ∼98%, while EEG provided a relative advantage for purely sensory tasks, though absolute accuracy remained more modest for both modalities. Critically, combining the two modalities together consistently matched or outperformed either one, used alone, across all conditions, with gains most pronounced for mixed vs sensory discrimination. Multi-subject generalization improved progressively with training-set size, rising to ∼88% with 15 training subjects for mixed classification, suggesting that subject-independent decoding of motor intent may be achievable when models are trained on a larger number of subjects. Taken together, these results establish that spinal ESG signals carry decodable information about peripheral stimulation that is complementary to and not redundant with cortical EEG. This finding supports a multilevel framework for decoding sensorimotor processing in humans and motivates the development of dual-modality brain–machine interfaces that leverage both cortical and spinal signals to improve the control of neurostimulation and assistive devices.

## Introduction

The somatosensory system provides an ideal model for investigating how sensory information is encoded and transformed along ascending neural pathways. In this system, afferent signals propagate through a sequence of well-defined anatomical relays, from peripheral nerves through the spinal cord and subcortical structures to primary and secondary somatosensory cortices (Abraira and Ginty, 2013; Coste et al., 2022; Wang et al., 2022). At each stage, neural populations transform the incoming signal through mechanisms such as temporal and spatial filtering, leading to progressively evolving sensory representations along the pathway (Gardner, 2001; Mountcastle, 2005; Pei et al., 2011). Peripheral electrical stimulation of mixed nerves, containing both sensory and motor fibers, provides a common experimental approach for studying the somatosensory systems. Indeed, electrical stimulation evokes highly reproducible responses with millisecond precision, whose amplitude and latency reflect the activity at successive levels of the somatosensory pathway. These responses, known as somatosensory evoked potentials (SEPs), have been extensively characterized using electrophysiological recordings. (Allison et al., 1991; Cruccu et al., 2008; Desmedt and Cheron, 1983; Rossini et al., 2015).

Studies of SEPs in humans have predominantly focused on cortical responses. These responses are commonly recorded using electroencephalography (EEG), which allows non-invasive measurement of cortical activity with high temporal resolution (Luck, 2005). Over several decades, researchers have characterized generators (Allison et al., 1991), scalp topographies (Mauguière et al., 1999), and latencies (Cruccu et al., 2008) of the major SEP components for stimulation of mixed nerves and individual digits. More recently, machine-learning approaches have extended this framework by showing that single-trial neural responses can, under certain conditions, contain sufficient information to classify stimulation site (Blankertz et al., 2011), and stimulus modality (Lin et al., 2024). In contrast to classical SEP analyses, which rely on trial averaging to isolate robust waveform components, decoding approaches assess whether patterns of activity in individual trials carry discriminative information about experimental conditions. Because such information may be distributed across channels, latencies, or spectral features, multivariate classifiers can reveal distinctions that remain invisible to conventional univariate analyses of average responses (Grootswagers et al., 2017). However, while these approaches establish EEG as a powerful tool for somatosensory classification (Craik et al., 2019; Saeidi et al., 2021), they primarily capture activity at the cortical endpoint of a multi-stage sensory pathway, leaving the contribution of earlier processing stages unresolved.

The spinal cord represents an ideal target for probing these earlier stages of somatosensory processing: it acts as a natural convergence point for afferent and efferent signals from multiple bilateral segments, making it an appealing recording site for capturing distributed sensorimotor activity with a single electrode array. Recent methodological advances now make it possible to record spinal activity non-invasively through high-density electrospinography (ESG), which enables spatially resolved recordings of spinal evoked activity using multichannel surface electrodes (Gabrieli et al., 2025; Mehra et al., 2025; Nierula et al., 2024). Although researchers detected spinal responses to peripheral stimulation at the body surface as early as the 1970s (Cracco, 1973; Shimoji et al., 1972), early ESG studies suffered from low spatial resolution and poor signal-to-noise ratios that limited their broader application. High-density ESG overcomes such limitations, by providing spatial information at the spinal level and by offering higher signal-to-noise ratio (SNR) due to signal combination from multiple electrodes (Gabrieli et al., 2025; Mehra et al., 2025; Nierula et al., 2024). In addition, combining high-density ESG with peripheral and cortical recordings opens the possibility for characterizing the full somatosensory pathway non-invasively and with high sensitivity. In particular, Nierula et al. (2024) showed that cervical and lumbar spinal responses to peripheral electrical stimulation can be reliably detected and spatially differentiated at the level of averaged evoked activity, establishing ESG as a viable tool for multilevel studies of human somatosensory processing. Such a platform could prove particularly relevant for clinical applications requiring the monitoring of sensorimotor processing at multiple levels, including mixed rehabilitation following spinal cord injury and somatosensory feedback restoration, where capturing distributed signals along the sensorimotor pathway could enhance decoding performance and reduce reliance on cortical recordings alone. However, it remains unclear whether spinal ESG signals contain decodable information at the single-trial level, and critically, whether they provide information that is complementary to, rather than redundant with, that captured by cortical EEG recordings.

In this work, we addressed this gap by testing whether spinal signals recorded with high-density ESG contain decodable information at the single-trial level and whether they provide complementary information to cortical EEG. We applied machine learning classification methods to EEG, ESG, and their combination to decode mixed (i.e., stimulation engaging both sensory and motor fibers), sensory, and mixed vs sensory stimulation conditions. We systematically compared classifier performance across modalities, identified optimal models for each task, and quantified the benefit of combining spinal and cortical signals. In addition, we assessed cross-subject generalization and characterized learning dynamics as a function of training data and training duration. This work aims to quantify the extent to which spinal ESG signals contribute to somatosensory decoding and whether their integration with EEG improves performance across tasks, laying the groundwork for multimodal neural interfaces that integrate neural activity at multiple levels of the sensorimotor pathway.

## Methods

### Dataset

The analyses presented in this study were conducted using a dataset of simultaneous electrospinography (ESG), electroencephalography (EEG) and electrocardiography (ECG) recordings previously published by Nierula et al. (Nierula et al., 2024) and publicly available on the OpenNeuro repository (“Somatosensory evoked potentials in the human spinal cord to mixed and sensory nerve stimulation - OpenNeuro,” n.d.). The dataset was acquired from 26 healthy participants and was originally designed to investigate non-invasive measurements of human spinal cord activity. In this work, we decided to include 17 of these participants (7 females, 23 ± 4 years old). The remaining nine participants were excluded due to corrupt or unreadable raw data files (5 participants), missing reference electrode recordings (1 participant), or excessive noise in ESG recordings (3 participants), as identified through inspection in the time and frequency domains.

### Experimental design

Here, we briefly summarize the experimental design. A detailed description of the experimental protocol and recording setup can be found in Nierula et al. (2024).

Electrophysiological signals were acquired non-invasively using TMS-compatible Ag/AgCl surface electrodes (Easycap GmbH, Germany). Recordings were performed using NeurOne Tesla amplifiers (Bittium Corporation, Finland) with a sampling rate of 10 kHz and an anti-aliasing filter between 0.16 and 2500 Hz. Electrode impedances were maintained below 10 kΩ during the recordings.

A schematic overview of the recording setup is shown in **Fig. 1A**. EEG activity was recorded using a cap-mounted montage of 39 electrodes positioned according to the international 10–10 system. Signals were referenced to the right mastoid and grounded at POz. ESG signals were recorded using a high-density surface electrode system consisting of 39 electrodes positioned over the posterior neck and trunk. Two dense electrode grids were centred over the cervical and lumbar spinal regions around anatomical target electrodes located at the sixth cervical vertebra (SC6) and the first lumbar vertebra (L1). Additional target electrodes were placed over SC1 and L4 to serve as anatomical landmarks. ESG signals were referenced to an electrode positioned over the sixth thoracic vertebra (T6). Two ventral electrodes placed in the supraglottic (AC) and supraumbilical (AL) regions were included to facilitate the extraction of spinal evoked potentials. ECG activity was also recorded using a bipolar configuration.

**Figure 1.**
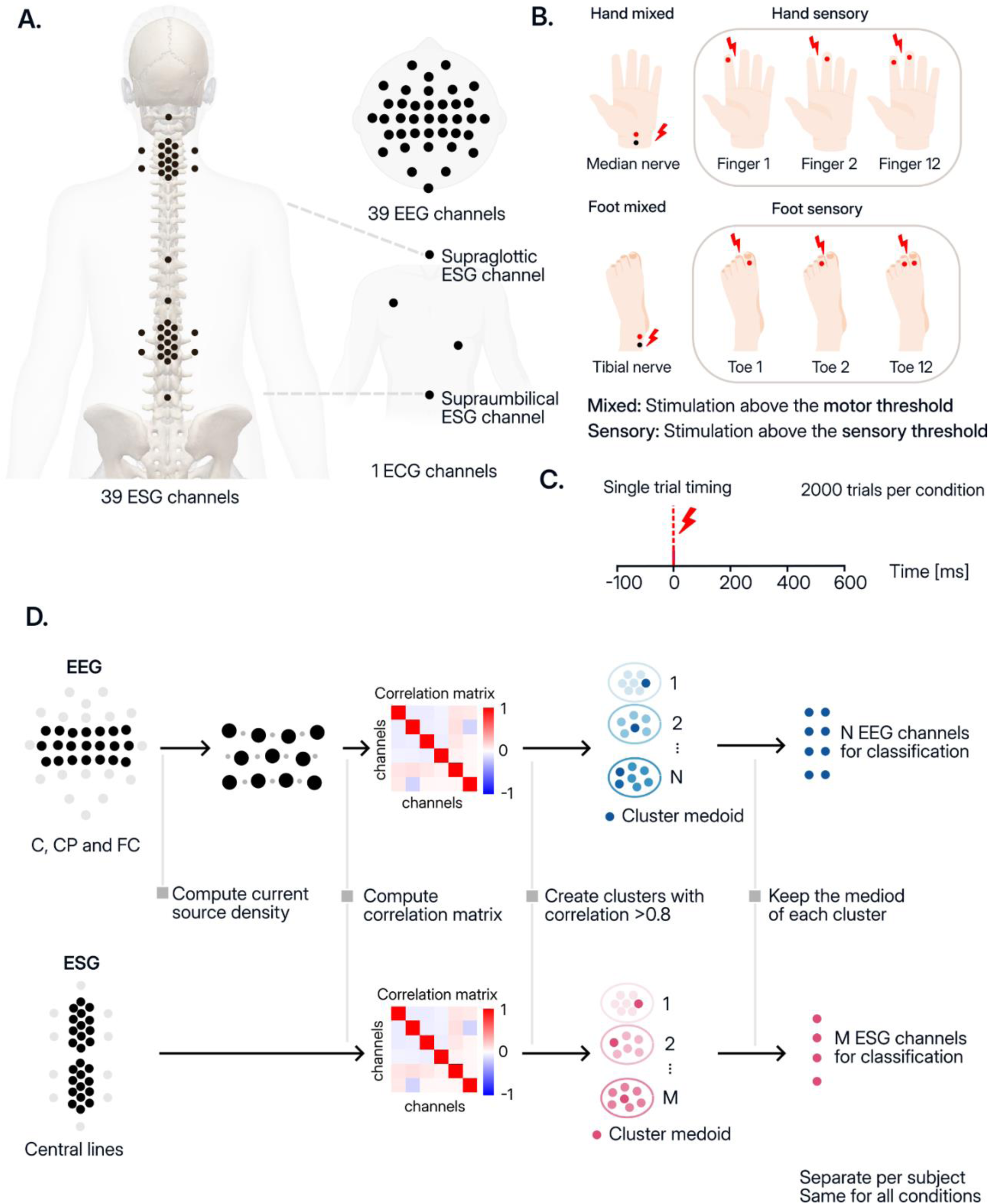
Experimental setup, stimulation conditions, and channel selection procedure. **A.** Placement of recording electrodes. Electroencephalography (EEG) was recorded from 39 scalp channels, electrospinography (ESG) from 39 electrodes positioned along the cervical and lumbar spinal levels, and electrocardiogram (ECG) from a single channel. **B.** Peripheral stimulation conditions. The dataset includes median nerve stimulation (*hand mixed* condition), stimulation of the index finger (*Finger 1*), middle finger (*Finger 2*), and simultaneous index–middle finger stimulation (*Finger 12*) (*hand sensory* condition); tibial nerve stimulation (*foot mixed* condition) and cutaneous stimulation of two toes delivered separately (*Toe 1, Toe 2*) and simultaneously (*Toe 12*) (*foot sensory* condition). Mixed conditions correspond to stimulation above mixed threshold, whereas sensory conditions correspond to stimulation above sensory threshold. **C.** Timing of a single trial. Stimulus onset occurred at 0 ms, and 2000 trials were acquired for each condition. **D.** Channel selection procedure. For EEG, current source density (CSD) was first computed from electrodes over central regions (C, CP, and FC). Pairwise channel correlations were then calculated, and channels were grouped into clusters with correlation > 0.8. One representative channel (cluster medoid) was retained from each cluster for classification. The same clustering procedure was applied to ESG channels recorded along the spinal midline. Channel selection was performed separately for each subject but identically across conditions.

Peripheral electrical stimulation was used to elicit somatosensory responses (**Fig. 1B**). Mixed nerve stimulation was delivered to the left median nerve at the wrist and to the left tibial nerve at the ankle using bipolar surface electrodes with an inter-electrode distance of 25 mm. Stimulation intensities were set just above the individual mixed threshold, which was defined as the intensity at which a participant’s thumb or first toe started to twitch (visually determined). Sensory stimulation targeted the index and middle fingers of the hand and the first and second toes of the foot using ring electrodes, with stimulation intensity set to three times the individual sensory detection threshold. Electrical pulses consisted of 0.2 ms square-wave stimuli delivered using constant-current stimulators (DS7A, Digitimer, UK). Each experimental session began with a 5-minute resting-state recording followed by stimulation blocks separated by short breaks. Mixed nerve blocks consisted of 2000 stimuli per condition. Sensory stimulation was delivered across four blocks per limb, with each individual digit condition (finger 1, finger 2, or both fingers; toe 1, toe 2, or both toes) presented 500 times per block, for a total of 2000 stimuli per condition. Stimuli were presented with a fixed inter-stimulus interval of 257 ms with an additional temporal jitter of ±20 ms.

### Data preprocessing

Unless otherwise stated, EEG, ESG, and ECG signals were preprocessed using a custom pipeline implemented in MNE-Python (version 1.10.2, Gramfort et al., 2013). For all recording modalities, preprocessing began with the removal of electrical stimulation artifacts occurring at stimulus onset. These artifacts were removed using interpolation based on piecewise cubic Hermite interpolating polynomials (PCHIP) (Higham, 1992). For ESG signals, interpolation windows were defined separately for cervical and lumbar channels based on visual inspection of the averaged responses. The largest cervical interpolation window extended from −5 ms to +6 ms relative to stimulus onset (11 ms total), while the largest lumbar interpolation window extended from −4 ms to +8 ms (12 ms total). For EEG and ECG signals, a fixed interpolation window of 5 ms was applied across participants. After artifact interpolation, EEG and ESG signals were preprocessed separately using different pipelines, as described below.

#### ESG preprocessing

ESG preprocessing followed the methodology described by Nierula et al. (2024). Cardiac artifacts in ESG recordings were removed using the simultaneously recorded ECG signal. R-peaks were detected using the FMRIB plug-in for the EEGLAB toolbox, run in MATLAB R2024a. Cardiac artifact removal was then performed using a channel-wise implementation of the Principal Component Analysis–Optimal Basis Set (PCA-OBS) method. Noisy channels were identified using power spectral density estimated with Welch’s method as implemented with MNE function. ESG data were re-referenced using an anterior electrode as reference, specifically the supraglottic electrode (AC) for median nerve stimulation and the supraumbilical electrode (AL) for tibial nerve stimulation, as this configuration has been shown to improve the extraction of spinal evoked potentials (Nierula et al., 2024). Data were band-pass filtered between 30 and 400 Hz with additional notch filtering at 50 Hz and its harmonics. Finally, preprocessed ESG data was segmented into epochs from −200 to 300 ms relative to stimulation onset, with baseline correction applied using the pre-stimulus interval (−100 to -10 ms).

#### EEG preprocessing

EEG data were downsampled to 1 kHz and visually inspected to identify noisy channels. Ocular artifacts were removed using independent component analysis (ICA) with the Picard algorithm (Ablin et al., 2018). Signals were then notch filtered at 50 Hz and its harmonics and band-pass filtered between 4 and 150 Hz using a finite impulse response (FIR) filter with a *firwin* design. Bad channels were interpolated using the MNE-Python *interpolate_bads()* function, which implements spherical spline interpolation (Perrin et al., 1989). Finally, the data were segmented into epochs from −200 to 300 ms relative to stimulation onset, and baseline correction was applied using the pre-stimulus interval (−100 to -10 ms).

### Decoding analysis

#### Channel selection

As a first step in the decoding pipeline, we applied a correlation-based channel selection procedure to reduce inter-channel redundancy and limit the dimensionality of the input data prior to classification (**Fig. 1D**). The procedure was performed independently for EEG and ESG signals and for each participant, but it was the same for all stimulation conditions.

A preliminary spatial restriction of channels was applied for both modalities. For EEG signals, we restricted the analysis to 21 channels over the FC, C, and CP regions, corresponding to sensorimotor cortical areas where somatosensory evoked responses are expected to be largest (Allison et al., 1991; Mauguière et al., 1999). Epochs were then re-referenced to the common average and transformed using current source density (CSD) via the MNE-Python *compute_current_source_density()* function. This transformation corresponds to the surface Laplacian of scalp potentials and reduces the effects of volume conduction while enhancing spatial resolution by emphasizing local cortical generators (Perrin et al., 1989). CSD was applied to EEG only, as scalp recordings are particularly susceptible to volume conduction and benefit from improved spatial specificity. For ESG recordings, CSD was not applied because the spinal electrode array was designed to capture distributed segmental spinal potentials along the rostrocaudal axis, thus preserving the native spatial and temporal morphology of these signals was prioritized over spatial sharpening. For ESG recordings, analysis was therefore restricted to 26 central spinal electrodes, including channels distributed along the cervical and lumbar segments and two adjacent parallel lines. Extreme lateral electrodes, brainstem electrodes, and anterior reference channels were excluded.

In addition, a supplementary analysis was conducted on a subset of five subjects to evaluate whether classification accuracy would improve for the *sensory* and *mixed vs. sensory* conditions when using a broader channel selection. In this analysis, all channels over the FC, C, and CP regions were included for EEG, and all spinal channels were retained except for extreme lateral electrodes, brainstem electrodes, and anterior reference channels.

For both modalities, epochs were cropped from 0 to 0.3 s after stimulus onset to focus the analysis on the stimulus-evoked response window. For each stimulation condition, a channel-by-channel Pearson correlation matrix was computed using epochs concatenated across trials and time samples. Channel pairs with an absolute correlation coefficient ≥ 0.8 were considered highly correlated (Mukaka, 2012). Only pairs that were consistently highly correlated across all stimulation conditions were retained, ensuring that the identified clusters reflected stable inter-channel relationships rather than condition-specific fluctuations. Highly correlated channel pairs were represented as a graph, where nodes correspond to channels and edges represent highly correlated pairs. Connected components of the graph defined clusters of redundant channels. For each cluster, the medoid channel, defined as the channel with the highest mean absolute correlation with all other channels in the cluster, was selected using the median mixed condition as a single reference condition to define subject-specific ESG channel groups. The resulting medoid channels were then applied across both median and tibial conditions to maintain consistent channel selection within participants and avoid redefining channels separately for each condition. Channels not belonging to any cluster were retained as singletons. The resulting channel sets for each participant are reported in **Supplementary Table 1** (ESG) and **Supplementary Table 2** (EEG).

#### Feature extraction

Decoding was performed using epochs cropped from 0 to 0.3 s after stimulus onset. For each participant and task, epochs were scaled using median-based normalization and flattened into single feature vectors by concatenating all selected channels and time points, as implemented through the MNE-Python *Scaler* and *Vectorizer* functions.

#### Single-subject classification tasks

Single-trial classification was performed to evaluate the discriminability of EEG and ESG responses between stimulation conditions at the level of individual epochs. We conducted decoding analyses independently for each participant using the reduced channel sets obtained from the correlation-based channel selection procedure described above.

Five classification tasks were considered:

1. **Mixed task**: The first task targeted suprathreshold mixed stimulation and aimed to discriminate stimulation location by distinguishing upper-limb (median mixed) from lower-limb (tibial mixed) conditions.
2. **Sensory tasks**: The second and third task targeted sensory stimulation and aimed to discriminate between digit-level conditions (finger 1 vs finger 2 and toe 1 vs toe 2, respectively).
3. **Mixed vs sensory tasks**: The fourth and fifth task addressed mixed–sensory differentiation by distinguishing mixed nerve stimulation from digit-level stimulation within the same limb (median mixed vs finger stimulation; tibial mixed vs toe stimulation). To avoid class imbalance due to pooling multiple digit-level sensory conditions, an equal number of epochs was randomly sampled from each sensory subclass before classifier training.

For each task, decoding was performed using EEG signals, ESG signals, and combined EEG&ESG signals. For ESG signals, channel selection also depended on the stimulation comparison. Cervical channels were used for finger-related comparisons, lumbar channels for toe-related comparisons, and both cervical and lumbar channels for hand vs foot comparisons.

#### Classifiers

We compared four supervised machine learning algorithms widely used in EEG decoding and brain–computer interface studies due to their robustness in high-dimensional neural data(Lotte et al., 2018, 2007). In particular, we considered Linear Discriminant Analysis (LDA), Support Vector Machines (SVM), and Multi-Layer Perceptron (MLP), implemented using scikit-learn version 1.7.2 (Pedregosa et al., 2012), and Extreme Gradient Boosting (XGBoost), implemented using the xgboost library (Chen and Guestrin, 2016).

#### Hyperparameter optimization

For each participant and task, the dataset was divided into training (80%) and test (20%) sets using stratified sampling with shuffling. Hyperparameter optimization was performed using the Optuna framework (Akiba et al., 2019) by maximizing mean classification accuracy through 10-fold stratified cross-validation on the training set. The search ranges for each classifier are reported in **Supplementary Table 3**. The best hyperparameter configuration was then used to refit the model on the full training data, and final classification performance was evaluated on the held-out test set. To further compare ESG and EEG performances across tasks, classification accuracies were summarized both by averaging performance across the four classifiers and by considering the best-performing classifier for each task.

#### Evaluation of classification performance as a function of trial number

To evaluate how the size of the training set influences decoding performance and to estimate the minimum number of trials required to achieve stable accuracy, an additional analysis was conducted by progressively varying the number of training epochs. This analysis was performed only for the ESG and ESG&EEG configurations, as our primary objective was to specifically assess the classification potential of this novel signal. For this reason, we focused exclusively on the mixed and mixed vs sensory tasks, since these were the conditions in which ESG demonstrated the highest decoding performance at the single-subject level (**Fig. 2**).

**Figure 2.**
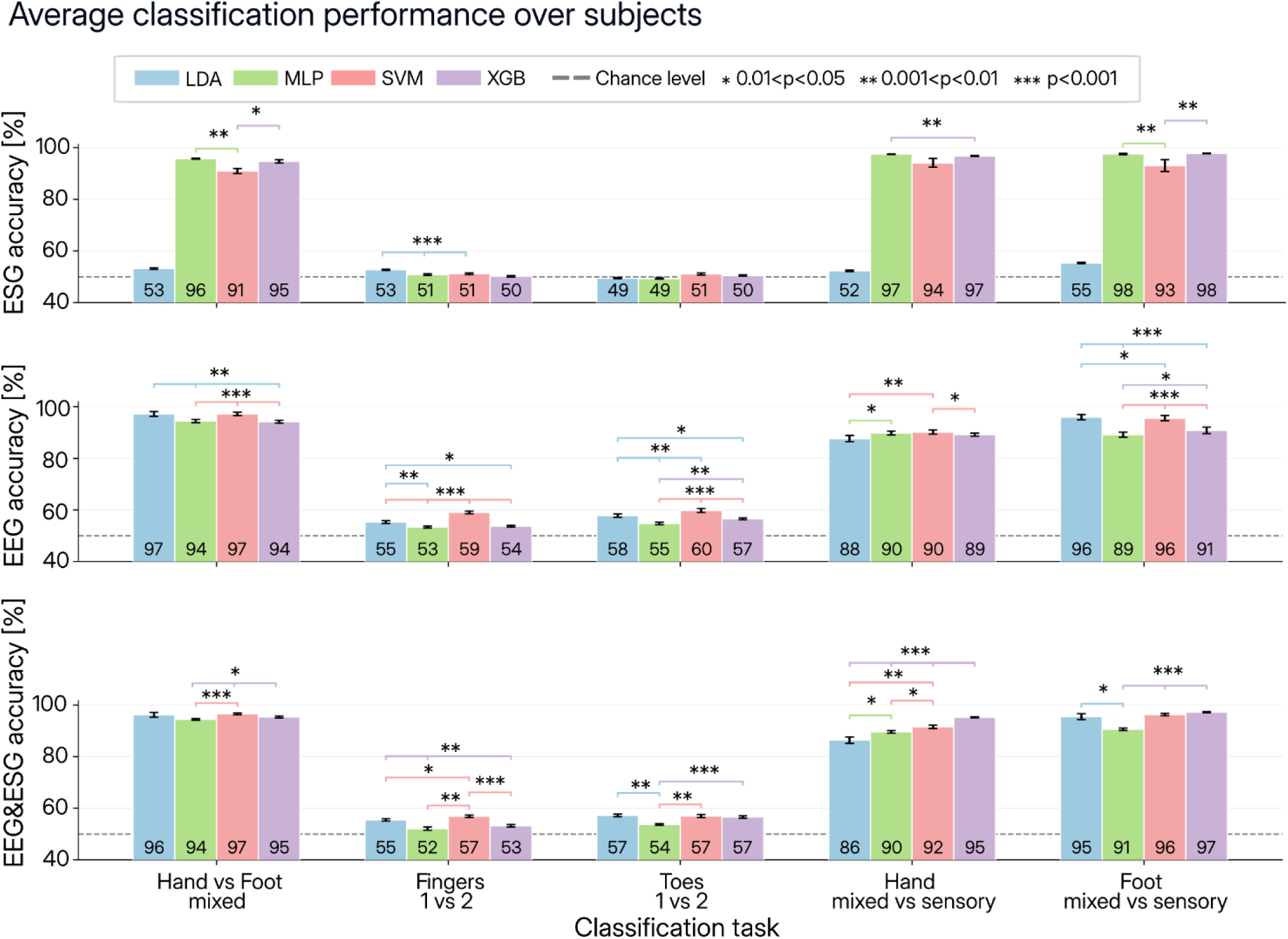
Classification performance across subjects and modalities. Average classification accuracy (%) across 17 subjects for five classification tasks using four classifiers: Linear Discriminant Analysis (LDA), Multi-Layer Perceptron (MLP), Support Vector Machine (SVM), and Extreme Gradient Boosting (XGB). Results are shown separately for ESG (top row), EEG (middle row), and the combined EEG&ESG modality (bottom row). Tasks include *Hand vs Foot mixed*, *Fingers 1 vs 2*, *Toes 1 vs 2*, *Hand mixed vs sensory*, and *Foot mixed vs sensory*. Bars represent the mean accuracy across subjects (also indicated at the base of each bar), with error bars indicating the standard error of the mean. The dashed horizontal line indicates chance level (50%). Asterisks denote statistically significant differences between classifiers (*\*\*\* = p<0.001, ** = 0.001<p<0.01, * = 0.01<p<0.05*). The color of the horizontal line beneath each asterisk corresponds to the classifier achieving the highest accuracy within that comparison.

For each participant, classification accuracy was evaluated as a function of the number of training epochs using the same preprocessed epochs and feature extraction procedure described in the previous sections. Training set sizes were varied from 100 to 3800 epochs (*N = 100, 300, 500, 1000, 1500, 2000, 2500, 3000, 3500, 3800*). For each value of N, a stratified random subset of N epochs was selected for training while all remaining epochs were used exclusively for testing. To account for variability due to random sampling, the selection of training epochs was repeated using ten different random seeds, resulting in ten independent training–test splits for each training set size.

Classification was performed using the best-performing model identified in the single-subject decoding analysis. For each participant and training size, classification accuracy was computed on the independent test set and averaged across repetitions. The minimum number of epochs required to reach stable performance was defined as the point beyond which further increases in training data produced negligible improvements in decoding accuracy, operationalized as an absolute increase of less than 3% between consecutive training steps (Śliwowski et al., 2023).

#### Multi-subject classification

To assess the generalizability of decoding performance across participants, we performed a multi-subject classification analysis, in which models were trained on data from a subset of participants and evaluated on held-out unseen participants. Multi-subject classification was performed only for the ESG and ESG&EEG configurations, as our primary objective was to specifically assess the classification potential of this novel signal. For this reason, we focused exclusively on the mixed and mixed vs sensory tasks, since these were the conditions in which ESG demonstrated the highest decoding performance at the single-subject level (**Figure 2**).

To ensure spatial consistency across participants, a common channel set was defined separately for each modality and task based on the frequency with which channels were retained after the correlation-based channel selection procedure. First, the average number of selected channels per participant was calculated for each modality-specific region, and then the channels with the highest selection frequency were retained up to that average number. For ESG, participants retained on average 1.24 ± 0.44 cervical channels and 1.24 ± 0.56 lumbar channels, resulting in SC6 and L1 as the most consistently selected cervical and lumbar channels, respectively; the four most frequently selected ESG channels overall were L1, SC6, S9, and S28. For EEG, participants retained an average of 7.76 channels, and the common EEG channel set therefore comprised the eight most frequently selected channels (C3, C4, C2, CP2, Cz, CPz, FC2, and CP1). The selected channels are shown in **Fig. 4A**.

Multi-subject decoding followed a leave-group-of-subjects-out strategy. For each selected task, models were trained on increasing numbers of participants (*N_subjects_* = 1, 3, 6, 9, 12, 15) and evaluated on the remaining held-out participants. For each training size, participants were randomly assigned to training and test sets using fixed random seeds, and results were averaged across ten repetitions with unique training-test splits. Hyperparameter optimization was performed on the training data only, using a subject-wise cross-validation strategy implemented via *GroupKFold* (n_splits = 5), ensuring that data from the same participant never appeared in both the training and validation folds.

### Statistical analysis

We performed statistical analyses using non-parametric tests, specifically the Wilcoxon signed-rank test (Pratt, 1959; Wilcoxon, 1945) and the Benjamini-Hochberg false discovery rate (FDR) correction for multiple comparisons (Benjamini and Hochberg, 1995), implemented via *scipy.stats.wilcoxon and statsmodels.stats.multitest.multipletests*, respectively.

#### Single-subject classification tasks

For the single-subject decoding analysis, classification accuracies for each participant, task, and classifier were compared against an empirical random baseline, estimated by simulating random binary predictions. Statistical significance was assessed using one-sided Wilcoxon signed-rank tests, testing whether observed accuracies exceeded this simulated chance level. False discovery rate FDR correction was applied separately for each signal modality (EEG, ESG, and EEG&ESG), with each family comprising 20 comparisons (5 classification tasks × 4 classifiers), using a corrected significance threshold of α = 0.05.

To identify the best-performing classifier for each task and signal modality, pairwise two-sided Wilcoxon signed-rank tests were applied to participant-level accuracies for all classifier pairs. FDR correction was applied globally across all pairwise comparisons. The classifier with the highest mean accuracy that was not significantly outperformed by any other classifier was selected as the best-performing classifier for each task. This classifier was subsequently used for the multi-subject classification and trial number analyses.

#### Comparing signal modalities

To compare decoding performance across signal modalities (EEG, ESG, and EEG&ESG), paired two-sided Wilcoxon signed-rank tests were applied to participant-level accuracies for each task, considering three pairwise comparisons: EEG&ESG vs EEG, EEG&ESG vs ESG, and ESG vs EEG. This analysis was performed both on accuracies averaged across the four classifiers and on accuracies obtained with the best-performing classifier per task. FDR correction was applied globally across all comparisons (15 tests: 5 tasks × 3 modality pairs), with a corrected significance threshold of α = 0.05.

#### Multi-subject classification

For the multi-subject decoding analysis, observed accuracies were compared against an empirical chance-level baseline derived from each test set’s class structure. Specifically, for each task, we simulated 2000 sets of random predictions by assigning class labels with equal probability and used the resulting accuracy distribution to estimate the performance expected by chance alone. One-sided Wilcoxon signed-rank tests were applied across the ten repetitions for each training size, testing whether observed accuracies exceeded the empirical baseline. FDR correction using the Benjamini–Hochberg procedure was applied within each task across the six training sizes (Ntrain = 1, 3, 6, 9, 12, 15), using a corrected significance threshold of α = 0.05.

## Results

### Single-subject decoding is robust for mixed tasks but limited for fine sensory discrimination

We first assessed overall decoding feasibility across tasks (mixed, sensory and mixed vs sensory) and modalities (ESG, EEG and combined EEG&ESG) (**Fig. 2**). Across all signal types, both mixed classification (*Hand vs Foot mixed*) and mixed vs sensory discrimination were robustly decodable, whereas fine sensory discrimination (*Fingers 1 vs 2 and Toes 1 vs 2)* remained substantially more challenging, with ESG performing near chance and EEG-based modalities showing only modest above-chance accuracy.

We compared classification performance across subjects using four classifiers (Linear Discriminant Analysis (LDA), Support Vector Machine (SVM), Multi-Layer Perceptron (MLP), and Extreme Gradient Boosting (XGB)) applied to ESG, EEG, and combined EEG&ESG signals (**Fig. 2**). Performance strongly depended on both task and modality. For *Hand vs Foot mixed* task, all modalities achieved high accuracy, with ESG reaching up to ∼96% (MLP), EEG ∼97% (LDA/SVM), and combined EEG&ESG ∼97% (SVM). In contrast, finer sensory discriminations (*Fingers 1 vs 2* and T*oes 1 vs 2*) proved substantially more challenging. For ESG, Fingers 1 vs 2 remained above chance for LDA, MLP, and SVM, but not XGB, while Toes 1 vs 2 exceeded chance only for SVM and XGB, with LDA and MLP remaining at chance level (**Supplementary Figure 2**). On the other hand, EEG and combined EEG&ESG achieved modest but above-chance accuracies (∼53–60%), with SVM and LDA generally outperforming XGB. For mixed versus sensory discrimination tasks, performance increased compared to purely sensory conditions. In the *Hand mixed vs sensory* condition, ESG reached ∼97% with nonlinear classifiers (MLP/XGB), while EEG achieved ∼88–90%, and combined EEG&ESG further improved performance up to ∼95% (XGB). A similar pattern was observed for the *Foot mixed vs sensory* task, where ESG and combined EEG&ESG achieved the highest accuracies (∼98% and ∼97%, respectively), outperforming EEG alone (∼89–96%). In summary, nonlinear classifiers (MLP and XGB) consistently yielded the highest performance in mixed-versus-sensory tasks for ESG and combined EEG&ESG signals, while LDA and SVM performed best for EEG-based decoding. For purely sensory discrimination tasks, EEG and combined EEG&ESG data achieved above-chance performance, with LDA and SVM showing similarly competitive accuracies. Based on these findings, we selected XGB for subsequent analyses of mixed and mixed-versus-sensory tasks, and LDA for sensory-only classifications, as it provided comparable performance to SVM while requiring approximately half training time.

To qualitatively assess whether strong decoding performance depended on dense electrode coverage, we compared classification accuracy obtained using all available channels versus the reduced correlation-based channel set in a representative subset of five participants (**Supplementary Fig. 3**). In this control analysis, the extended configuration included all EEG channels over the FC, C, and CP regions and all spinal channels except for extreme lateral electrodes, brainstem electrodes, and anterior reference channels. Across ESG, EEG, and combined EEG&ESG modalities, reduced channel configurations generally preserved or improved classification performance. For example, in the Hand mixed vs sensory task, ESG accuracy increased from 83% to 97%, EEG from 64–75% to 89–91%, and combined EEG&ESG from 81% to 89–95% when using selected rather than extended channel configurations. Similarly, in the Foot mixed vs sensory task, selected channels achieved 98% ESG accuracy and 97–98% combined EEG&ESG accuracy while matching or outperforming extended-channel analyses.

### EEG and combined EEG&ESG outperform ESG in sensory decoding

We next compared classification performance across signal modalities (EEG, ESG, and combined EEG&ESG), first averaging across classifiers and then selecting the best-performing classifier for each task (**Fig. 3**). When averaged across classifiers (**Fig. 3A**), modality-dependent differences were evident across task types. For the *Hand vs foot mixed* mixed task, EEG and combined EEG&ESG achieved the highest accuracies (∼96%), both outperforming ESG (∼84%). In contrast, for sensory discrimination (*Fingers 1 vs 2* and *Toes 1 vs 2*), performance remained close to chance for all modalities, although EEG and EEG&ESG (∼54–57%) significantly outperformed ESG (∼50–51%, all p < 0.001). For mixed-versus-sensory tasks, accuracy increased substantially across all modalities. In the *Hand mixed vs sensory* condition, EEG (∼89%) and EEG&ESG (∼91%) outperformed ESG (∼85%). Similarly, for *Foot mixed vs sensory*, combined EEG&ESG achieved the highest accuracy (∼95%), followed by EEG (∼93%) and ESG (∼86%). When selecting the best-performing classifier for each task (**Fig. 3B**), ESG and EEG&ESG performance improved. For the *Hand vs foot mixed* task, all modalities reached high accuracy (∼94–95%). For mixed-versus-sensory tasks, ESG and combined EEG&ESG achieved the highest performance, reaching up to ∼97–98%.

**Figure 3.**
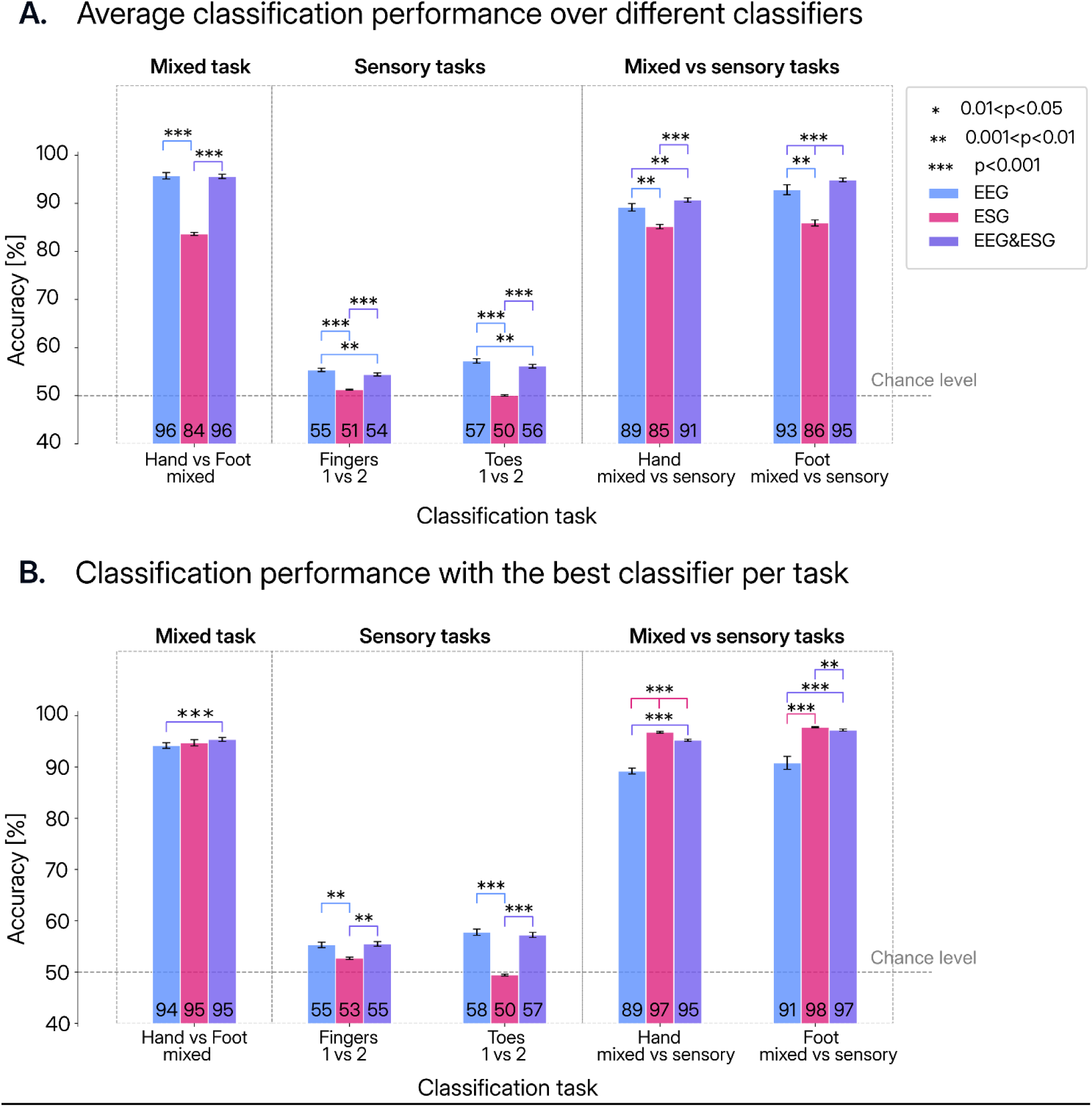
Average classification performance across subjects and modalities. **A.** Average classification accuracy across 17 subjects for EEG, ESG, and the combined EEG&ESG modalities, averaged across all classifiers: Linear Discriminant Analysis (LDA), Support Vector Machine (SVM), Multi-Layer Perceptron (MLP), and Extreme Gradient Boosting (XGB). Tasks are grouped into mixed (*Hand vs foot mixed*), sensory (*Fingers 1 vs 2*, *Toes 1 vs 2*), and mixed vs sensory (*Hand mixed vs sensory*, *Foot mixed vs sensory*) conditions. Bars represent mean accuracy across subjects with standard error of the mean. The dashed horizontal line indicates chance level (50%). Numbers at the base of each bar indicate the mean accuracy. **B.** Classification accuracy when selecting the best-performing classifier for each task. Linear Discriminant Analysis (LDA) yielded the best performance for sensory discrimination tasks, whereas Extreme Gradient Boosting (XGB) performed best for mixed and mixed-versus-sensory tasks. Asterisks denote statistically significant differences between modalities (*\*\*\* = p<0.001, ** = 0.001<p<0.01, * = 0.01<p<0.05*). The color of the horizontal line beneath each asterisk corresponds to the classifier achieving the highest accuracy within that comparison.

### Decoding performance plateaus rapidly with increasing training data

We next quantified how classification performance evolved with training epochs across subjects (**Fig. 5**). For the *Hand vs Foot mixed* task, accuracy increased rapidly and reached near-ceiling levels for both modalities, with mean performance approaching ∼95–100% within the first ∼500–1000 epochs for ESG and combined EEG&ESG signals. In contrast, mixed vs sensory discrimination tasks showed slower and more gradual improvements. For *Hand mixed vs sensory*, ESG accuracy increased from ∼60–70% at early epochs to ∼85–90% at plateau, while the combined EEG&ESG modality achieved higher final performance (∼88–92%). Similarly, for *Foot mixed vs sensory*, ESG performance went from ∼55–65% to ∼85–90%, whereas EEG&ESG reached ∼90–95%. Across all tasks, combining EEG with ESG consistently improved both final accuracy (by ∼3–7%) and convergence speed compared to ESG alone. Median plateau epochs, indicated by dashed vertical lines and summarized by violin plots, were typically around 1500 epochs, with earlier stabilization for *Hand vs Foot mixed* (1000 epochs). Notably, inter-subject variability was minimal for the binary mixed task but substantially larger for mixed vs sensory discrimination, as reflected by the wider spread of individual curves and plateau distributions.

### Multi-subject generalization is strong for mixed decoding and improves with combined EEG&ESG

We performed a multi-subject classification analysis to assess the generalizability of ESG-based decoding performance across participants. We selected a common set of channels for all subjects (**Fig. 4A**), and we trained the classifiers on an increasing number of subjects, while being evaluated on the remaining participants. For *Hand vs Foot mixed* task (**Fig. 4B**), classification accuracy increased steadily with the number of training subjects for both ESG and the combined EEG&ESG modality. Using a single training subject, accuracy was ∼74% for ESG and ∼73% for EEG&ESG. Performance improved progressively as more subjects were included, reaching ∼86% (ESG) and ∼88% (EEG&ESG) when trained on 15 subjects. Notably, all conditions performed significantly above chance level. For the *Hand mixed vs sensory* discrimination (**Fig. 4C**), a different pattern emerged. ESG-based classification remained close to chance across all training-set sizes (∼48–53%). In contrast, the combined EEG&ESG modality showed clear improvements with additional training data, increasing from ∼53% with one training subject to ∼66% when trained on 15 subjects. Finally, the *Foot mixed vs sensory* task (**Fig. 4D**), both ESG and EEG&ESG achieved above-chance performance. Accuracy increased with the number of training subjects, reaching ∼62% for ESG and ∼74% for EEG&ESG at the largest training set.

**Figure 4.**
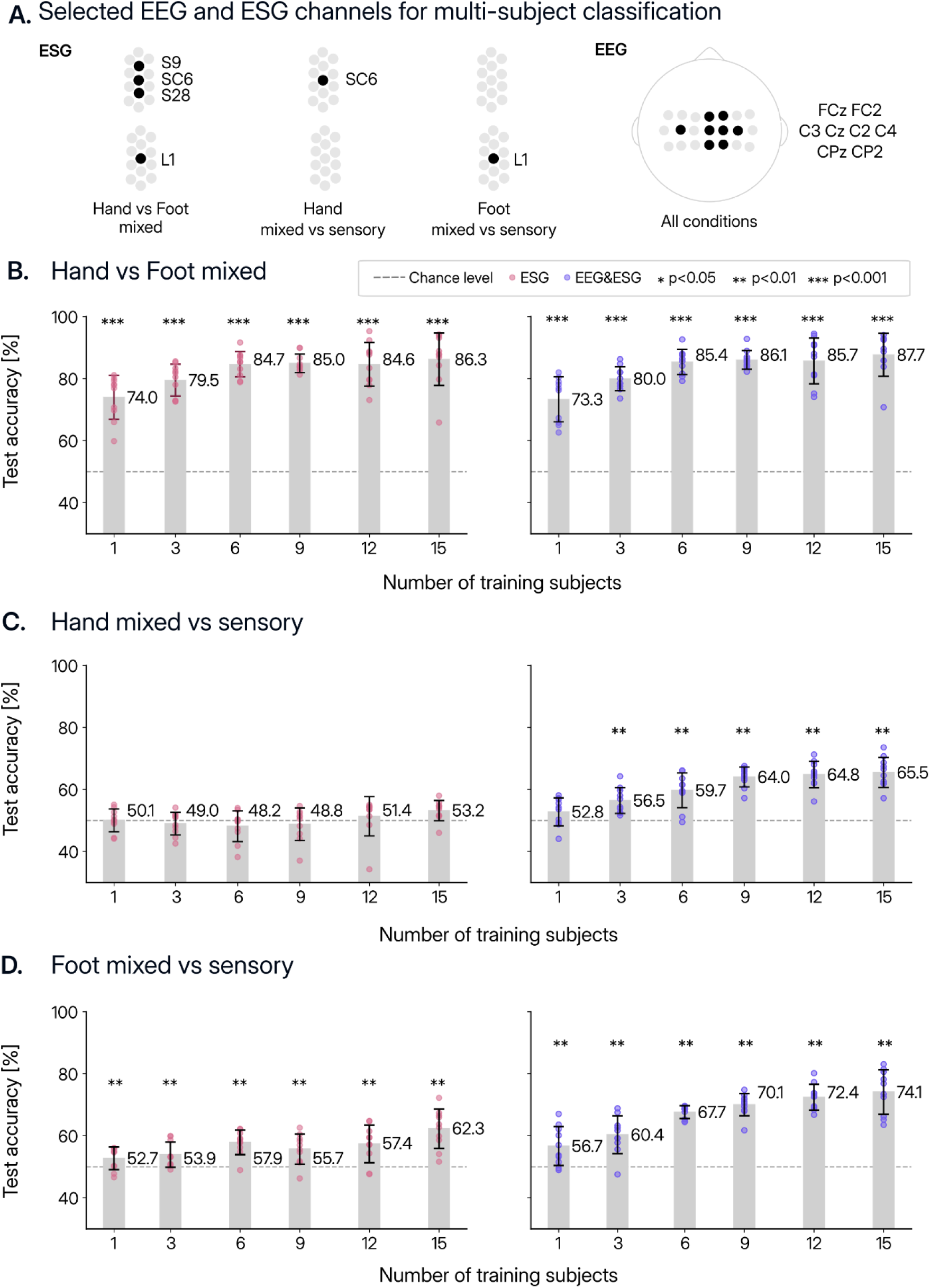
Multi-subject classification performance using ESG and combined EEG&ESG signals. **A.** Selected ESG and EEG channels used for multi-subject classification. Channels were identified by averaging cluster assignments from subject-specific clustering and retaining those most consistently represented across participants. **B-D** Classification accuracy as a function of the number of training subjects for three conditions in which ESG data was consistently above chance: **B.** *Hand vs foot mixed*, **C.** *hand mixed vs sensory*, and **D.** *foot mixed vs sensory*. Results are shown for ESG alone (left panels) and combined EEG&ESG signals (right panels). For each number of training subjects, classification was repeated 10 times using different train–test splits, with the remaining subjects serving as the test set. Dots represent individual repetitions, while bars indicate the mean accuracy across repetitions with standard deviation. Numerical values above bars denote mean accuracy (%). The dashed horizontal line indicates chance level (50%). Asterisks indicate statistical significance above chance level (*\*\*\* = p<0.001, ** = 0.001<p<0.01, * = 0.01<p<0.05*).

## Discussions

This study provides, to our knowledge, the first systematic investigation of electrospinography (ESG) as a tool for single-trial neural decoding, and the first direct comparison of spinal and cortical (EEG) signals within a unified multimodal decoding framework. Three principal findings emerge from our results. First, ESG alone achieves highly accurate decoding of mixed-related conditions, reaching up to ∼96% accuracy for *Hand vs Foot mixed* classification in single-subject analyses (**Fig. 3**) and ∼86% in multi-subject generalization with 15 training participants (**Fig. 4**). Second, EEG provides modest but above-chance sensory decoding (∼54–60%, **Fig. 3**), complementing rather than duplicating the information carried by ESG, that performs at chance level for fine sensory discriminations. Third, multimodal fusion of EEG and ESG performed comparably to or better than the best single modality in single-subject decoding, while in cross-subject generalization it consistently improved performance relative to ESG alone, reaching ∼88% for mixed and ∼66–74% for mixed vs sensory tasks with the largest training sets (**Fig. 4**).

ESG has been recently used to characterize spinal cord physiology, including lower limb movement (Steele et al., 2024), responses to peripheral stimulation (Gabrieli et al., 2025; Nierula et al., 2024) and patterns of neural activity and corticospinal coherence during pincer-grip (Mehra et al., 2025), but its use for single-trial classification has not previously been demonstrated. Our results show that ESG supports highly accurate decoding in mixed stimulation conditions, even when using only a small number of channels (up to four per subject, **Supplementary Table 1**). One possible explanation is that mixed stimulation engages greater peripheral input and larger recruited neural populations, potentially producing larger and more spatially consistent ESG deflections that enhance decodability (Buzsáki et al., 2012). However, the specific physiological factors underlying this advantage (such as stimulation intensity, number of recruited fibers, or other protocol-dependent features) cannot be disentangled from the present design.

By contrast, ESG decoding of fine sensory discriminations (*Fingers 1 vs 2*, *Toes 1 vs 2*) remained at or near chance level across all classifiers (**Fig. 2**). This probably indicates that the somatotopic organization underlying fine tactile discrimination is either poorly preserved in the spinal field potential recorded at the body surface or falls below the signal-to-noise threshold accessible to non-invasive ESG (Farina et al., 2014; Nierula et al., 2024). Skin-surface recordings necessarily integrate activity across multiple spinal segments and are attenuated by intervening tissue, limiting spatial resolution (Michel and Brunet, 2019). Invasive approaches may partially overcome these constraints. For instance, epidural recordings have been shown to capture sensory information through modulations in theta and beta band activity (Shukla et al., 2024), indicating that such information is present at the spinal level but may not be detectable with current non-invasive methods. In addition, ESG recordings are susceptible to noise from surrounding muscles and other physiological sources (Nierula et al., 2024; Steele et al., 2024), further limiting sensitivity to fine-grained sensory signals. Future work combining high-density ESG arrays, improved acquisition hardware, and source-separation methods tailored to spinal cord geometry may help recover finer somatotopic structure and enhance sensory decoding performance.

EEG&ESG showed the most consistent advantage, especially in mixed-vs-sensory discrimination, confirming complementarity rather than substitutability of one of the two signals. This multimodal advantage is most compellingly demonstrated in the cross-subject generalization analysis, where the practical stakes are highest. For the *Hand vs Foot mixed* task, accuracy increased from ∼73% with a single training subject to ∼88% with 15 subjects for EEG&ESG, compared to ∼74% and ∼86% respectively for ESG alone. The more striking pattern emerged for mixed vs sensory discrimination. For the *Hand mixed vs sensory* task, ESG-based cross-subject decoding remained near chance across all training-set sizes (∼48–53%), while EEG&ESG improved steadily from ∼53% with one training subject to ∼66% with 15. For *Foot mixed vs sensory*, ESG reached ∼62% and EEG&ESG ∼74% at the largest training set. These results show that multimodal fusion qualitatively enables generalization in conditions where the spinal signal alone is uninformative across participants. Combining a modality that reflects activity closer to the effector and that is anatomically more constrained than cortical signals is conceptually analogous to EEG–fMRI fusion approaches in cognitive neuroscience, where complementary temporal and spatial information are jointly exploited (Huster et al., 2012), as well as EEG–EMG coupling, which links cortical dynamics to peripheral motor output (Lee et al., 2025; Ortega-Auriol et al., 2023; von Carlowitz-Ghori et al., 2014). In our case, this complementarity operates along the sensorimotor neuroaxis, with cortical and spinal signals capturing distinct but synergistic aspects of the underlying processes (Capogrosso et al., 2016; Rowald et al., 2022; Wagner et al., 2018).

One contributing factor to the high performance in mixed-vs-sensory tasks may be differences in SNR across stimulation conditions, as double finger or toe stimulation is known to produce somatosensory evoked potentials with higher SNR compared to single-digit stimulation (Nierula et al., 2024). When contrasting mixed nerve stimulation with sensory-only conditions, SNR may therefore not be fully matched across classes, potentially inflating class separability. However, SNR differences alone cannot fully account for the results. If they were the primary driver, both ESG and EEG&ESG would be expected to benefit equally. Instead, in cross-subject generalization, ESG remained near chance while EEG&ESG improved systematically, pointing to complementary cortical information as a key contributor to decoding performance.

All experimental conditions relied on peripheral electrical stimulation, which elicits strong, time-locked afferent volleys with inherently high signal-to-noise ratio. Whether comparable decoding performance can be achieved during voluntary movement or passive sensory processing remains an open question. Mixed artefacts represent a more serious confound for ESG than for EEG, and their impact on single-trial classification in naturalistic paradigms has not yet been quantified (Muthukumaraswamy, 2013; Sburlea and Müller-Putz, 2018). Although Steele et al. (2023) demonstrated that ESG signals can characterize voluntary movements, single-trial decoding of movement versus rest or between distinct movements was not attempted. Whether ESG-based decoding extends to voluntary motor conditions therefore remains to be established.

Beyond these limitations, our findings position the combined EEG&ESG platform as a promising candidate for closed-loop BCI applications requiring bidirectional communication, such as motor rehabilitation following spinal cord injury (Rowald et al., 2022; Wagner et al., 2018) and somatosensory feedback systems (Flesher et al., 2016; Raspopovic et al., 2014). A key practical advantage is that high decoding accuracy was achieved using a reduced electrode montage. Despite this limited spatial sampling, performance remained high for mixed and mixed vs sensory tasks, was comparable to current state-of-the-art BCI systems (Roy et al., 2019; Schirrmeister et al., 2017). Importantly, control analyses in a subset of subjects demonstrated that extending the channel set to include all EEG electrodes over FC, C, and CP regions and all spinal channels (excluding extreme lateral, brainstem, and anterior reference electrodes) did not systematically improve performance (**Supplementary Fig. 3**). In several cases, reduced channel configurations even yielded higher accuracies, indicating that the correlation-based selection procedure effectively captures the most informative signals while reducing redundancy.

Such reduced-channel configurations are particularly relevant for clinical deployment, as they minimize setup time, hardware complexity, and user burden, factors that remain major barriers to real-world BCI adoption (Lotte et al., 2018; McFarland and Wolpaw, 2017). In addition, the strong cross-subject generalization observed for combined EEG&ESG suggests that multimodal approaches may reduce the need for extensive subject-specific calibration, thereby lowering the barrier to practical use. Accuracy approached ∼88% with 15 training subjects (**Fig. 4**), indicating that subject-independent decoding may be achievable for certain conditions. Although performance for mixed vs sensory tasks remained more modest, the consistent improvement with increasing training-set size suggests that larger datasets could further enhance generalization. Finally, analysis of learning dynamics (**Fig. 5**) showed that performance typically plateaued within ∼1000–1500 epochs, with faster convergence observed for mixed tasks. This is consistent with previous ESG findings indicating that stable evoked responses can be obtained with a few hundred trials (Nierula et al., 2024). Reducing the number of required training epochs has direct implications for shortening calibration time and improving usability in applied settings. Together, these results point toward a new class of multimodal BCIs that leverage distributed neural representations across the sensorimotor neuroaxis to enable more robust and generalizable human–machine interfaces.

**Figure 5.**
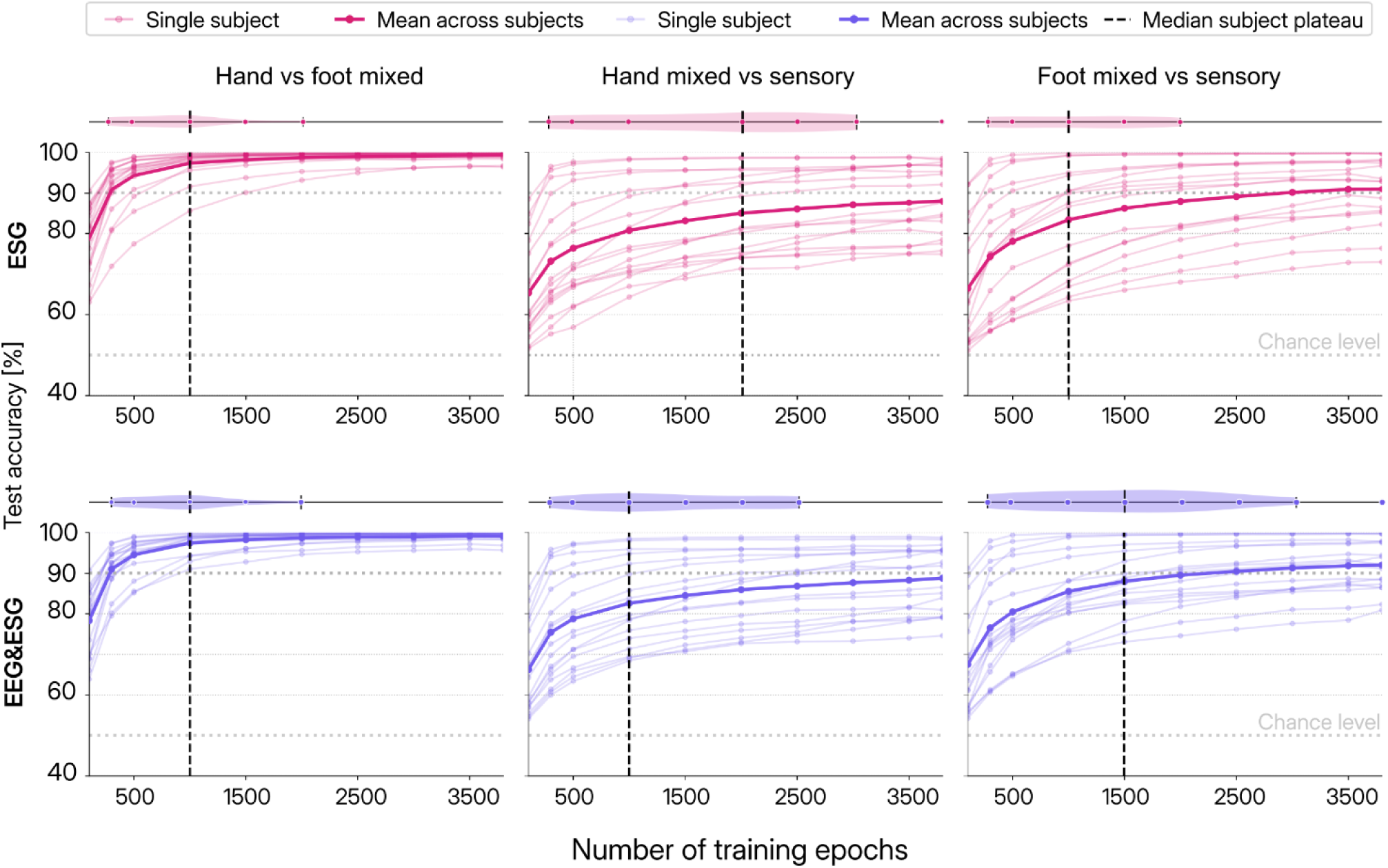
Learning curves and plateau analysis across subjects for ESG and combined EEG&ESG signals. Classification accuracy as a function of the number of training epochs for three tasks: *Hand vs Foot (mixed)*, *Hand (mixed vs sensory)*, and *Foot (mixed vs sensory)*. Results are shown for ESG (top row) and combined EEG&ESG signals (bottom row). Thin lines represent individual subject performance, while thick lines indicate the mean accuracy across subjects. The dashed vertical line marks the median number of training epochs at which subjects reach a performance plateau. The horizntal dashed line indicates chance level (50%). Violin plots shown above each panel summarize the distribution of plateau points across subjects, illustrating variability in the number of epochs required to reach stable performance. Performance increases rapidly with training epochs for all tasks, with the fastest convergence observed for *Hand vs Foot (mixed)*.

## Data and Code availability

The dataset analyzed in this study was previously published by Nierula et al. and is publicly available through the OpenNeuro repository (“Somatosensory evoked potentials in the human spinal cord to mixed and sensory nerve stimulation - OpenNeuro,” n.d.). The custom code used for data preprocessing, decoding analyses, and statistical evaluation is available from the corresponding author upon reasonable request.

## Ethics statement

This study analyzed publicly available de-identified data from the OpenNeuro repository originally published by (Nierula et al., 2024). The original study was approved by the Ethics Committee at the Medical Faculty of the University of Leipzig, and all participants provided written informed consent. No additional ethical approval was required for the present secondary analysis.

## Author contributions

**L.T.** Conceptualization, Methodology, Software, Investigation, Formal analysis, Data curation, Writing-Original Draft, Writing – Review & Editing, Visualization; **G.C.** Conceptualization, Methodology, Software, Investigation, Formal analysis, Data curation, Writing-Original Draft, Writing – Review & Editing, Visualization; **A.M.** Software, Formal analysis, Data curation, Writing – Review & Editing; **B.N.** Methodology, Software, Data curation, Writing – Review & Editing, Visualization **D.E.** Resources, Writing – Review & Editing; **F.A.** Resources, Methodology, Writing – Review & Editing; **S.I.** Resources, Writing – Review & Editing; **F.E.** Methodology, Resources, Writing – Review & Editing; **F.A.** Conceptualization, Methodology, Resources, Visualization, Writing – Review & Editing, Supervision, Project administration; **S.M.** Conceptualization, Resources, Supervision, Project administration, Funding acquisition, Writing – Review & Editing.

## References

Ablin, P., Cardoso, J.-F., Gramfort, A., 2018. Faster Independent Component Analysis by Preconditioning With Hessian Approximations. IEEE Trans. Signal Process. 66, 4040–4049. 10.1109/TSP.2018.2844203

Abraira, V.E., Ginty, D.D., 2013. The sensory neurons of touch. Neuron 79, 618–639. 10.1016/j.neuron.2013.07.051

Akiba, T., Sano, S., Yanase, T., Ohta, T., Koyama, M., 2019. Optuna: A Next-generation Hyperparameter Optimization Framework. 10.48550/arXiv.1907.10902

Allison, T., McCarthy, G., Wood, C.C., Jones, S.J., 1991. Potentials evoked in human and monkey cerebral cortex by stimulation of the median nerve. A review of scalp and intracranial recordings. Brain J. Neurol. 114 (Pt 6), 2465–2503. 10.1093/brain/114.6.2465

Benjamini, Y., Hochberg, Y., 1995. Controlling the False Discovery Rate: A Practical and Powerful Approach to Multiple Testing. J. R. Stat. Soc. Ser. B Methodol. 57, 289–300.

Blankertz, B., Lemm, S., Treder, M., Haufe, S., Müller, K.-R., 2011. Single-trial analysis and classification of ERP components -- A tutorial 814–825. 10.1016/j.neuroimage.2010.06.048

Buzsáki, G., Anastassiou, C.A., Koch, C., 2012. The origin of extracellular fields and currents — EEG, ECoG, LFP and spikes. Nat. Rev. Neurosci. 13, 407–420. 10.1038/nrn3241

Capogrosso, M., Milekovic, T., Borton, D., Wagner, F., Moraud, E.M., Mignardot, J.-B., Buse, N., Gandar, J., Barraud, Q., Xing, D., Rey, E., Duis, S., Jianzhong, Y., Ko, W.K.D., Li, Q., Detemple, P., Denison, T., Micera, S., Bezard, E., Bloch, J., Courtine, G., 2016. A brain–spine interface alleviating gait deficits after spinal cord injury in primates. Nature 539, 284–288. 10.1038/nature20118

Chen, T., Guestrin, C., 2016. XGBoost: A Scalable Tree Boosting System, in: Proceedings of the 22nd ACM SIGKDD International Conference on Knowledge Discovery and Data Mining. pp. 785–794. 10.1145/2939672.2939785

Controlling the False Discovery Rate: A Practical and Powerful Approach to Multiple Testing | Journal of the Royal Statistical Society Series B: Statistical Methodology | Oxford Academic [WWW Document], n.d. URL https://academic.oup.com/jrsssb/article/57/1/289/7035855?login=false (accessed 3.16.26).

Coste, C.A., William, L., Fonseca, L., Hiairrassary, A., Andreu, D., Geffrier, A., Teissier, J., Fattal, C., Guiraud, D., 2022. Activating effective functional hand movements in individuals with complete tetraplegia through neural stimulation. Sci. Rep. 12, 16189. 10.1038/s41598-022-19906-x

Cracco, R.Q., 1973. Spinal evoked response: Peripheral nerve stimulation in man. Electroencephalogr. Clin. Neurophysiol. 35, 379–386. 10.1016/0013-4694(73)90195-8

Craik, A., He, Y., Contreras-Vidal, J.L., 2019. Deep learning for electroencephalogram (EEG) classification tasks: a review. J. Neural Eng. 16, 031001. 10.1088/1741-2552/ab0ab5

Cruccu, G., Aminoff, M.J., Curio, G., Guerit, J.M., Kakigi, R., Mauguiere, F., Rossini, P.M., Treede, R.-D., Garcia-Larrea, L., 2008. Recommendations for the clinical use of somatosensory-evoked potentials. Clin. Neurophysiol. 119, 1705–1719. 10.1016/j.clinph.2008.03.016

Desmedt, J.E., Cheron, G., 1983. Spinal and far-field components of human somatosensory evoked potentials to posterior tibial nerve stimulation analysed with oesophageal derivations and non-cephalic reference recording. Electroencephalogr. Clin. Neurophysiol. 56, 635–651. 10.1016/0013-4694(83)90031-7

Farina, D., Jiang, N., Rehbaum, H., Holobar, A., Graimann, B., Dietl, H., Aszmann, O.C., 2014. The extraction of neural information from the surface EMG for the control of upper-limb prostheses: emerging avenues and challenges. IEEE Trans. Neural Syst. Rehabil. Eng. Publ. IEEE Eng. Med. Biol. Soc. 22, 797–809. 10.1109/TNSRE.2014.2305111

Flesher, S.N., Collinger, J.L., Foldes, S.T., Weiss, J.M., Downey, J.E., Tyler-Kabara, E.C., Bensmaia, S.J., Schwartz, A.B., Boninger, M.L., Gaunt, R.A., 2016. Intracortical microstimulation of human somatosensory cortex. Sci. Transl. Med. 8, 361ra141–361ra141. 10.1126/scitranslmed.aaf8083

Gabrieli, G., Somervail, R., Mouraux, A., Leandri, M., Haggard, P., Iannetti, G.D., 2025. Electrical Spinal Imaging: A noninvasive, high-resolution approach that enables electrophysiological mapping of the human spinal cord. PLOS Biol. 23, e3003116. 10.1371/journal.pbio.3003116

Gardner, E.P., 2001. Touch, in: Encyclopedia of Life Sciences. John Wiley & Sons, Ltd. 10.1038/npg.els.0000219

Gramfort, A., Luessi, M., Larson, E., Engemann, D.A., Strohmeier, D., Brodbeck, C., Goj, R., Jas, M., Brooks, T., Parkkonen, L., Hämäläinen, M., 2013. MEG and EEG data analysis with MNE-Python. Front. Neurosci. 7, 267. 10.3389/fnins.2013.00267

Higham, D.J., 1992. Monotonic piecewise cubic interpolation, with applications to ODE plotting. J. Comput. Appl. Math. 39, 287–294. 10.1016/0377-0427(92)90205-C

Huster, R.J., Debener, S., Eichele, T., Herrmann, C.S., 2012. Methods for simultaneous EEG-fMRI: an introductory review. J. Neurosci. Off. J. Soc. Neurosci. 32, 6053–6060. 10.1523/JNEUROSCI.0447-12.2012

Lee, H.-T., Shim, M., Liu, X., Cheon, H.-R., Kim, S.-G., Han, C.-H., Hwang, H.-J., 2025. A review of hybrid EEG-based multimodal human–computer interfaces using deep learning: applications, advances, and challenges. Biomed. Eng. Lett. 15, 587–618. 10.1007/s13534-025-00469-5

Lin, H.-Y., He, C., Su, C.-H., Hope Pan, L.-L., Hsiao, F.-J., Wu, Y.-T., Wang, Y.-F., Wang, S.-J., Ko, L.-W., 2024. Decoding Human Somatosensory Sensitivity Through Resting EEG and Behavioral Analysis: A Multimodal Fusion Approach. IEEE Trans. Neural Syst. Rehabil. Eng. Publ. IEEE Eng. Med. Biol. Soc. 32, 3310–3319. 10.1109/TNSRE.2024.3434353

Lotte, F., Bougrain, L., Cichocki, A., Clerc, M., Congedo, M., Rakotomamonjy, A., Yger, F., 2018. A review of classification algorithms for EEG-based brain-computer interfaces: a 10 year update. J. Neural Eng. 15, 031005. 10.1088/1741-2552/aab2f2

Lotte, F., Congedo, M., Lécuyer, A., Lamarche, F., Arnaldi, B., 2007. A review of classification algorithms for EEG-based brain-computer interfaces. J. Neural Eng. 4, R1–R13. 10.1088/1741-2560/4/2/R01

Luck, S.J. (Steven J., 2005. An introduction to the event-related potential technique. Cambridge, Mass.: MIT Press.

Mauguière, F., Allison, T., Babiloni, C., Buchner, H., Eisen, A.A., Goodin, D.S., Jones, S.J., Kakigi, R., Matsuoka, S., Nuwer, M., Rossini, P.M., Shibasaki, H., 1999. Somatosensory evoked potentials. The International Federation of Clinical Neurophysiology. Electroencephalogr. Clin. Neurophysiol. Suppl. 52, 79–90.

McFarland, D.J., Wolpaw, J.R., 2017. EEG-Based Brain-Computer Interfaces. Curr. Opin. Biomed. Eng. 4, 194–200. 10.1016/j.cobme.2017.11.004

Mehra, P., Bista, S., Metzger, M., Giglia, E.R., Plaitano, S., Nash, L., Domhnaill, É.M., Bede, P., Muthuraman, M., Hardiman, O., Lowery, M., Nasseroleslami, B., 2025. Spinal electrophysiology reveals frequency-specific spatial patterns of neural activity and corticospinal coherence during pincer-grip. 10.1101/2025.11.11.687864

Michel, C.M., Brunet, D., 2019. EEG Source Imaging: A Practical Review of the Analysis Steps. Front. Neurol. 10, 325. 10.3389/fneur.2019.00325

Mountcastle, V.B., 2005. The Sensory Hand: Neural Mechanisms of Somatic Sensation. Harvard University Press. 10.2307/j.ctv23dxd9k

Mukaka, M.M., 2012. Statistics corner: A guide to appropriate use of correlation coefficient in medical research. Malawi Med. J. J. Med. Assoc. Malawi 24, 69–71.

Muthukumaraswamy, S.D., 2013. High-frequency brain activity and muscle artifacts in MEG/EEG: a review and recommendations. Front. Hum. Neurosci. 7, 138. 10.3389/fnhum.2013.00138

Nierula, B., Stephani, T., Bailey, E., Kaptan, M., Pohle, L.-M.G., Horn, U., Mouraux, A., Maess, B., Villringer, A., Curio, G., Nikulin, V.V., Eippert, F., 2024. A multichannel electrophysiological approach to noninvasively and precisely record human spinal cord activity. PLOS Biol. 22, e3002828. 10.1371/journal.pbio.3002828

Ortega-Auriol, P., Byblow, W.D., Besier, T., McMorland, A.J.C., 2023. Muscle synergies are associated with intermuscular coherence and cortico-synergy coherence in an isometric upper limb task. Exp. Brain Res. 241, 2627–2643. 10.1007/s00221-023-06706-6

Pedregosa, F., Varoquaux, G., Gramfort, A., Michel, V., Thirion, B., Grisel, O., Blondel, M., Prettenhofer, P., Weiss, R., Dubourg, V., Vanderplas, J., Passos, A., Cournapeau, D., Brucher, M., Perrot, M., Duchesnay, E., Louppe, G., 2012. Scikit-learn: Machine Learning in Python. J. Mach. Learn. Res. 12.

Pei, Y.C., Hsiao, S.S., Craig, J.C., Bensmaia, S.J., 2011. Neural mechanisms of tactile motion integration in somatosensory cortex. Neuron 69, 536–547. 10.1016/j.neuron.2010.12.033

Perrin, F., Pernier, J., Bertrand, O., Echallier, J.F., 1989. Spherical splines for scalp potential and current density mapping. Electroencephalogr. Clin. Neurophysiol. 72, 184–187. 10.1016/0013-4694(89)90180-6

Pratt, J.W., 1959. Remarks on Zeros and Ties in the Wilcoxon Signed Rank Procedures. J. Am. Stat. Assoc. 54, 655–667. 10.1080/01621459.1959.10501526

Raspopovic, S., Capogrosso, M., Petrini, F.M., Bonizzato, M., Rigosa, J., Di Pino, G., Carpaneto, J., Controzzi, M., Boretius, T., Fernandez, E., Granata, G., Oddo, C.M., Citi, L., Ciancio, A.L., Cipriani, C., Carrozza, M.C., Jensen, W., Guglielmelli, E., Stieglitz, T., Rossini, P.M., Micera, S., 2014. Restoring natural sensory feedback in real-time bidirectional hand prostheses. Sci. Transl. Med. 6, 222ra19. 10.1126/scitranslmed.3006820

Rossini, P.M., Burke, D., Chen, R., Cohen, L.G., Daskalakis, Z., Di Iorio, R., Di Lazzaro, V., Ferreri, F., Fitzgerald, P.B., George, M.S., Hallett, M., Lefaucheur, J.P., Langguth, B., Matsumoto, H., Miniussi, C., Nitsche, M.A., Pascual-Leone, A., Paulus, W., Rossi, S., Rothwell, J.C., Siebner, H.R., Ugawa, Y., Walsh, V., Ziemann, U., 2015. Non-invasive electrical and magnetic stimulation of the brain, spinal cord, roots and peripheral nerves: Basic principles and procedures for routine clinical and research application. An updated report from an I.F.C.N. Committee. Clin. Neurophysiol. Off. J. Int. Fed. Clin. Neurophysiol. 126, 1071–1107. 10.1016/j.clinph.2015.02.001

Rowald, A., Komi, S., Demesmaeker, R., Baaklini, E., Hernandez-Charpak, S.D., Paoles, E., Montanaro, H., Cassara, A., Becce, F., Lloyd, B., Newton, T., Ravier, J., Kinany, N., D’Ercole, M., Paley, A., Hankov, N., Varescon, C., McCracken, L., Vat, M., Caban, M., Watrin, A., Jacquet, C., Bole-Feysot, L., Harte, C., Lorach, H., Galvez, A., Tschopp, M., Herrmann, N., Wacker, M., Geernaert, L., Fodor, I., Radevich, V., Van Den Keybus, K., Eberle, G., Pralong, E., Roulet, M., Ledoux, J.-B., Fornari, E., Mandija, S., Mattera, L., Martuzzi, R., Nazarian, B., Benkler, S., Callegari, S., Greiner, N., Fuhrer, B., Froeling, M., Buse, N., Denison, T., Buschman, R., Wende, C., Ganty, D., Bakker, J., Delattre, V., Lambert, H., Minassian, K., van den Berg, C.A.T., Kavounoudias, A., Micera, S., Van De Ville, D., Barraud, Q., Kurt, E., Kuster, N., Neufeld, E., Capogrosso, M., Asboth, L., Wagner, F.B., Bloch, J., Courtine, G., 2022. Rowald2022 - Activity-dependent spinal cord neuromodulation rapidly restores trunk and leg motor functions after complete paralysis. Nat. Med. 28, 260–271. 10.1038/s41591-021-01663-5

Roy, Y., Banville, H., Albuquerque, I., Gramfort, A., Falk, T.H., Faubert, J., 2019. Deep learning-based electroencephalography analysis: a systematic review. J. Neural Eng. 16, 051001. 10.1088/1741-2552/ab260c

Saeidi, M., Karwowski, W., Farahani, F.V., Fiok, K., Taiar, R., Hancock, P.A., Al-Juaid, A., 2021. Neural Decoding of EEG Signals with Machine Learning: A Systematic Review. Brain Sci. 11, 1525. 10.3390/brainsci11111525

Sburlea, A.I., Müller-Putz, G.R., 2018. Exploring representations of human grasping in neural, muscle and kinematic signals. Sci. Rep. 8, 16669. 10.1038/s41598-018-35018-x

Schirrmeister, R.T., Springenberg, J.T., Fiederer, L.D.J., Glasstetter, M., Eggensperger, K., Tangermann, M., Hutter, F., Burgard, W., Ball, T., 2017. Deep learning with convolutional neural networks for EEG decoding and visualization. Hum. Brain Mapp. 38, 5391–5420. 10.1002/hbm.23730

Shimoji, K., Kano, T., Higashi, H., Morioka, T., Henschel, E.O., 1972. Evoked spinal electrograms recorded from epidural space in man. J. Appl. Physiol. 33, 468–471. 10.1152/jappl.1972.33.4.468

Shukla, P.D., Burke, J.F., Kunwar, N., Presbrey, K., Balakid, J., Yaroshinsky, M., Louie, K., Jacques, L., Shirvalkar, P., Wang, D.D., 2024. Human Cervical Epidural Spinal Electrogram Topographically Maps Distinct Volitional Movements. J. Neurosci. 44. 10.1523/JNEUROSCI.2258-23.2024

Śliwowski, M., Martin, M., Souloumiac, A., Blanchart, P., Aksenova, T., 2023. Impact of dataset size and long-term ECoG-based BCI usage on deep learning decoders performance. Front. Hum. Neurosci. 17, 1111645. 10.3389/fnhum.2023.1111645

Somatosensory evoked potentials in the human spinal cord to mixed and sensory nerve stimulation - OpenNeuro [WWW Document], n.d. URL https://openneuro.org/datasets/ds004389/versions/1.0.0 (accessed 4.24.26).

Steele, A.G., Faraji, A.H., Contreras-Vidal, J.L., 2024. Electrospinography for non-invasively recording spinal sensorimotor networks in humans. J. Neural Eng. 20, 066043. 10.1088/1741-2552/ad1782

von Carlowitz-Ghori, K., Bayraktaroglu, Z., Hohlefeld, F.U., Losch, F., Curio, G., Nikulin, V.V., 2014. Corticomuscular coherence in acute and chronic stroke. Clin. Neurophysiol. Off. J. Int. Fed. Clin. Neurophysiol. 125, 1182–1191. 10.1016/j.clinph.2013.11.006

Wagner, F.B., Mignardot, J.-B., Le Goff-Mignardot, C.G., Demesmaeker, R., Komi, S., Capogrosso, M., Rowald, A., Seáñez, I., Caban, M., Pirondini, E., Vat, M., McCracken, L.A., Heimgartner, R., Fodor, I., Watrin, A., Seguin, P., Paoles, E., Van Den Keybus, K., Eberle, G., Schurch, B., Pralong, E., Becce, F., Prior, J., Buse, N., Buschman, R., Neufeld, E., Kuster, N., Carda, S., von Zitzewitz, J., Delattre, V., Denison, T., Lambert, H., Minassian, K., Bloch, J., Courtine, G., 2018. Targeted neurotechnology restores walking in humans with spinal cord injury. Nature 563, 65–71. 10.1038/s41586-018-0649-2

Wang, L.-H., Ding, W.-Q., Sun, Y.-G., 2022. Spinal ascending pathways for somatosensory information processing. Trends Neurosci. 45, 594–607. 10.1016/j.tins.2022.05.005

Wilcoxon, F., 1945. Individual Comparisons by Ranking Methods. Biom. Bull. 1, 80–83. 10.2307/3001968

