## Supplementary Material for "Decoding peripheral stimulation from cortical and spinal recordings reveals complementary sensorimotor information"

This supplementary material provides additional methodological details and supporting results, including electrode configurations for ESG and EEG recordings, statistical analyses of classification performance across modalities, tasks, and classifiers, and comparisons between full and reduced channel sets based on correlation-driven selection. Supplementary tables report subject-specific channel selection outcomes for ESG and EEG, as well as hyperparameter search ranges for all classifiers.

### Supplementary Figure 1

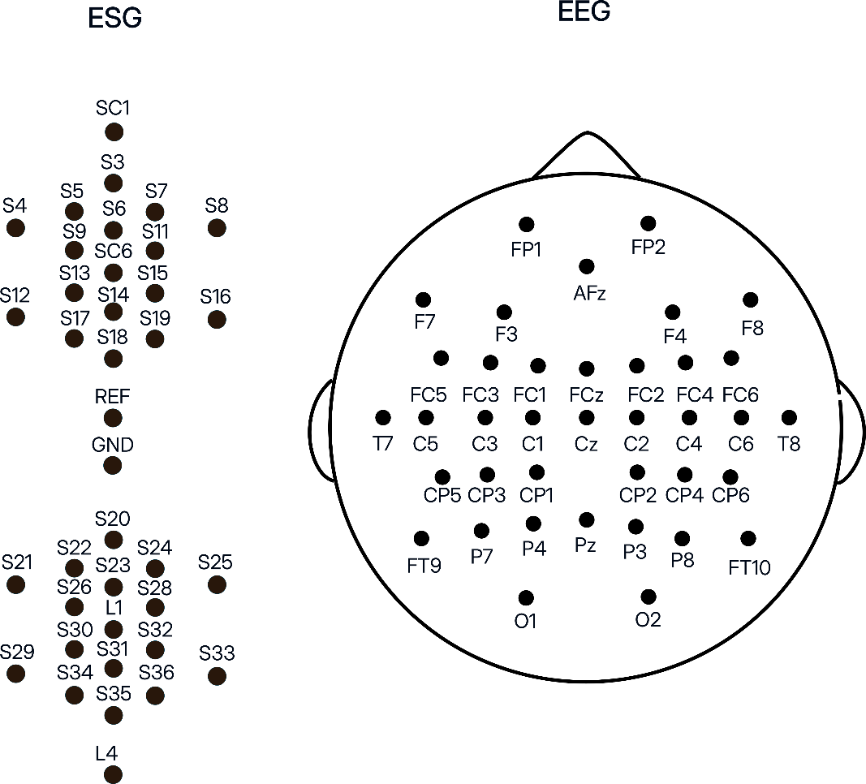

**Supplementary figure 1. Electrode configuration for ESG and EEG recordings.**
Schematic representation of electrode placement for electrospinography (ESG; left) and electroencephalography (EEG; right). ESG electrodes were distributed along the cervical, thoracic, and lumbar regions of the spinal cord, covering segments from upper cervical (e.g., SC1) to lower lumbar levels (e.g., L4), with reference (REF) and ground (GND) electrodes indicated. EEG electrodes were positioned according to the international 10–10 system, covering frontal, central, parietal, temporal, and occipital regions.

### Supplementary Figure 2

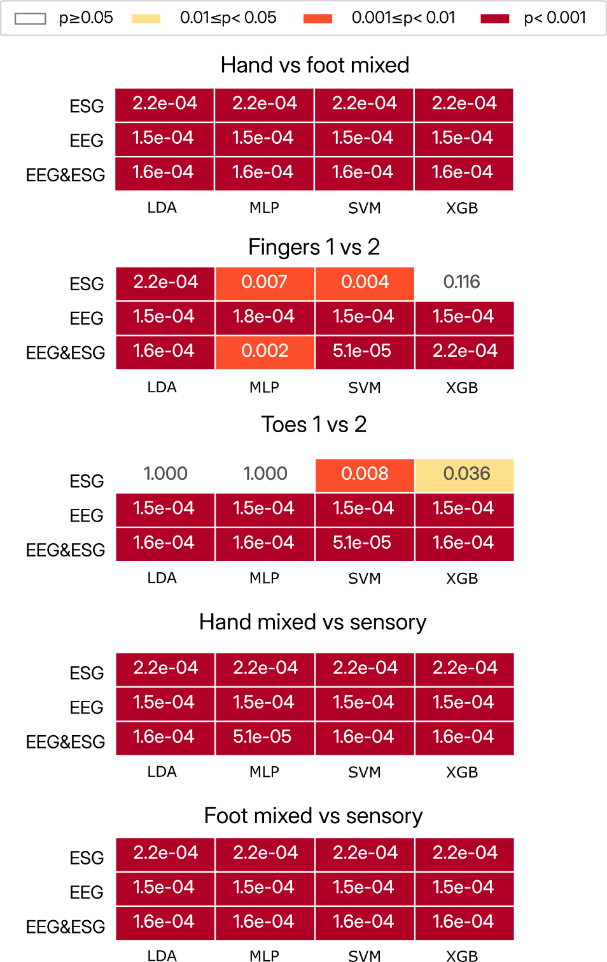

**Supplementary Figure 2. Statistical significance of classification performance relative to empirical chance level across signal modalities and tasks.** Heatmaps show false discovery rate–corrected p-values from one-sided Wilcoxon signed-rank tests assessing whether single-subject classification accuracies exceeded an empirical random baseline derived from simulated binary predictions. Results are shown separately for spinal recordings, cortical recordings, and the combined modality across five classification tasks (hand vs foot mixed, fingers 1 vs 2, toes 1 vs 2, hand mixed vs sensory, and foot mixed vs sensory) and four classifiers (linear discriminant analysis (LDA), multilayer perceptron (MLP), support vector machine (SVM), and extreme gradient boosting (XGB)). False discovery rate correction was applied separately within each modality across 20 comparisons (5 tasks × 4 classifiers; corrected α = 0.05). Cell colors indicate significance level (white: p ≥ 0.05; yellow: 0.01 ≤ p < 0.05; orange: 0.001 ≤ p < 0.01; dark red: p < 0.001), and numeric values within each cell report the corrected p-value.

### Supplementary Figure 3

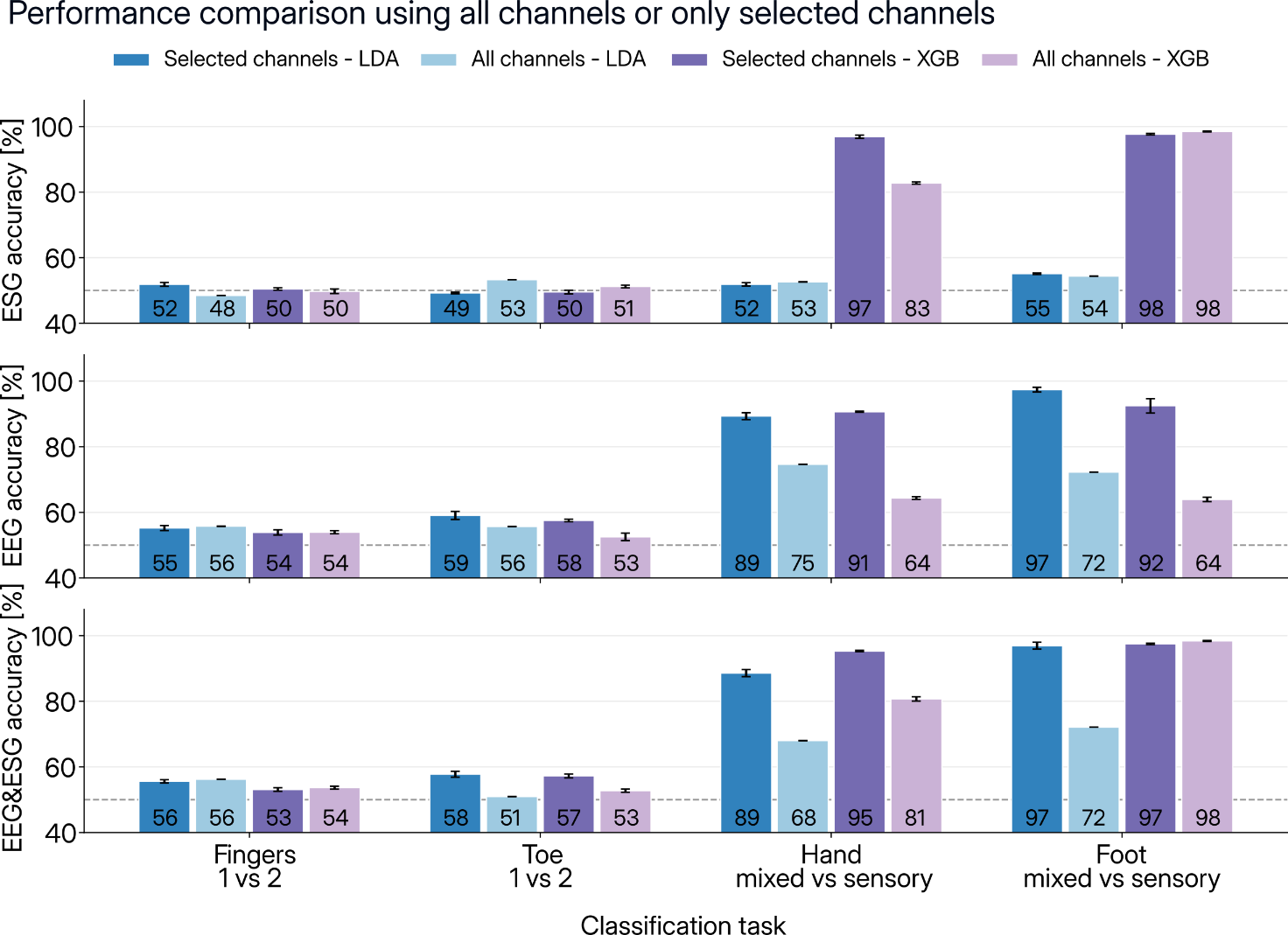

**Supplementary Figure 3.** **Representative comparison of classification performance using all available channels versus correlation-based selected channels in five example subjects.** Average classification accuracy (%) across subjects for ESG (top), EEG (middle), and combined EEG&ESG (bottom) signals across four classification tasks: Fingers 1 vs 2, Toes 1 vs 2, Hand mixed vs sensory, and Foot mixed vs sensory. For each task, performance is shown for Linear Discriminant Analysis (LDA) and Extreme Gradient Boosting (XGB) using either the reduced channel set obtained through the correlation-based channel selection procedure (*Selected channels*) or an extended channel set (*All channels*). In the extended configuration, EEG analyses included all channels over the FC, C, and CP regions, while ESG analyses included all spinal channels except for extreme lateral electrodes, brainstem electrodes, and anterior reference channels. Bars represent mean accuracy across subjects, with error bars indicating the standard error of the mean. Numerical values at the base of each bar indicate mean accuracy (%).

### Supplementary Table 1

| **Subject** | **Initial ESG channels (n)** | **Clusters (n)** | **Medoids** | **Singletons** | **Final ESG channels** |
| --- | --- | --- | --- | --- | --- |
| A | 26 | 2 | L1, S9 | S7, S31 | L1, S9, S7, S31 |
| B | 26 | 3 | L1, S17, S26 | – | L1, S17, S26 |
| C | 26 | 3 | L1, S11, S18 | – | L1, S11, S18 |
| D | 26 | 2 | L1, S9 | S24 | L1, S9, S24 |
| E | 26 | 2 | L1, S11 | – | L1, S11 |
| F | 26 | 2 | L1, SC6 | – | L1, SC6 |
| G | 26 | 4 | S15, S26, S28, S35 | – | S15, S26, S28, S35 |
| H | 26 | 2 | S15, S28 | – | S15, S28 |
| I | 26 | 2 | L1, SC6 | – | L1, SC6 |
| J | 26 | 2 | S32, SC6 | – | S32, SC6 |
| K | 26 | 2 | S28, SC6 | S3 | S28, SC6, S3 |
| L | 26 | 2 | L1, S13 | – | L1, S13 |
| M | 26 | 2 | S34, S9 | – | S34, S9 |
| N | 26 | 2 | L1, S11 | – | L1, S11 |
| O | 26 | 2 | S24, SC6 | – | S24, SC6 |
| P | 26 | 2 | S28, S9 | – | S28, S9 |
| Q | 26 | 2 | L1, SC6 | – | L1, SC6 |

**Table 1.** Summary of correlation-based channel selection results for ESG channels for all subjects.

### Supplementary Table 2

| **Subject** | **Initial EEG channels (n)** | **Clusters (n)** | **Medoids** | **Singletons** | **Final EEG channels** |
| --- | --- | --- | --- | --- | --- |
| A | 21 | 4 | C3, C4, CP1, FCz | FC2, CP2, FC5, C2 | C3, C4, FC2, CP1, CP2, FC5, FCz, C2 |
| B | 21 | 4 | C3, C4, CP1, FC1 | Cz, FC2, CP2, C1, CPz | C3, C4, Cz, FC1, FC2, CP1, CP2, C1, CPz |
| C | 21 | 3 | C3, C4, FC1 | Cz, CP1, CP2, FC5, C2, CPz | C3, C4, Cz, FC1, CP1, CP2, FC5, C2, CPz |
| D | 21 | 3 | C3, C4, CPz | Cz, FC2, CP1, FCz, C2 | C3, C4, Cz, FC2, CP1, FCz, C2, CPz |
| E | 21 | 3 | C3, C4, CPz | Cz, FC5 | C3, C4, Cz, FC5, CPz |
| F | 21 | 2 | C3, C4 | Cz, FC1, CP1, CP2, FCz, C1, CPz | C3, C4, Cz, FC1, CP1, CP2, FCz, C1, CPz |
| G | 21 | 4 | C2, C3, C4, Cz | FC2 | C3, C4, Cz, FC2, C2 |
| H | 21 | 4 | C3, C4, CPz, FCz | FC2, C2 | C3, C4, FC2, FCz, C2, CPz |
| I | 21 | 3 | C4, CP3, FC3 | Cz, FC1, FC2, CP2, FCz, C1, C2, CPz | C4, Cz, FC1, FC2, CP2, FCz, C1, C2, FC3, CP3, CPz |
| J | 21 | 5 | C1, C3, CP6, CPz, FCz | FC5, FC6, C2, FC4 | C3, FC5, FC6, CP6, FCz, C1, C2, FC4, CPz |
| K | 21 | 2 | C3, C4 | Cz, CP1, CP2, FCz, C1, C2, CPz | C3, C4, Cz, CP1, CP2, FCz, C1, C2, CPz |
| L | 21 | 2 | C5, CPz | CP2, FC5 | CP2, FC5, C5, CPz |
| M | 21 | 1 | C4 | C1 | C4, C1 |
| N | 21 | 3 | C3, C4, FC1 | Cz, FC2, CP1, CP2, C2, CPz | C3, C4, Cz, FC1, FC2, CP1, CP2, C2, CPz |
| O | 21 | 6 | C1, C3, C4, CP1, FC1, FC5 | CP2, C2 | C3, C4, FC1, CP1, CP2, FC5, C1, C2 |
| P | 21 | 3 | C3, C6, CP2 | Cz, FC1, FC2, CP1, FCz, C1, C2, FC4, C6 | C3, Cz, FC1, FC2, CP1, FCz, C1, C2, FC4, C6 |
| Q | 21 | 3 | C1, C3, C4 | Cz, FC1, FC2, CP2, FCz, C2, CPz | C3, C4, Cz, FC1, FC2, CP2, FCz, C1, C2, CPz |

**Table 2.** Summary of correlation-based channel selection results for EEG channels for all subjects.

### Supplementary Table 3

| **Classifier** | **Hyperparameter** | **Search range** |
| --- | --- | --- |
| **LDA** | Solver | {svd, lsqr, eigen} |
| **SVM** | Kernel | {linear, rbf} |
|  | C | (1e−3, 1e3), log scale |
|  | γ (rbf only) | {scale, auto} |
| **MLP** | Hidden layer sizes | {(50,), (100,), (100, 50), (150,)} |
|  | α (L2 regularization) | (1e−6, 1e−2), log scale |
|  | Learning rate init | (1e−4, 1e−2), log scale |
|  | Activation | {relu, tanh} |
| **XGBoost** | n_estimators | (100, 600) |
|  | max_depth | (2, 8) |
|  | Learning rate | (1e−3, 0.3), log scale |
|  | Subsample | (0.5, 1.0) |
|  | colsample_bytree | (0.5, 1.0) |
|  | λ (L2 regularization) | (1e−3, 10.0), log scale |
|  | min_child_weight | (1e−2, 10.0), log scale |

**Table 3** Hyperparameter search ranges for each classifier.
